# Dual targeting of transferrin receptor and CD98hc enhances brain exposure of large molecules

**DOI:** 10.1101/2025.03.24.645085

**Authors:** Robert C. Wells, Padma Akkapeddi, Darren Chan, David Joy, Roni Chau, Johann Chow, Arash Moshkforoush, Gerald Maxwell Cherf, Ellen Y.T. Chien, Allisa Clemens, Michelle E. Pizzo, Ahlam N. Qerqez, Tiffany Tran, Isabel Becerra, Jason C. Dugas, Joseph Duque, Timothy K. Earr, David Huynh, Do Jin Kim, Amy Wing-Sze Leung, Eric Liang, Hoang N. Nguyen, Yaneth Robles-Colmenares, Holly Kane, Elysia Roche, Patricia Sacayon, Kaitlin Xa, Hilda Solanoy, Mark S. Dennis, Joseph W. Lewcock, Ryan J. Watts, Mihalis S. Kariolis, Robert G. Thorne, Christopher M. Koth, Meredith E. K. Calvert, Y. Joy Yu Zuchero, Kylie S. Chew

## Abstract

Targeting proteins highly expressed at the blood-brain barrier, including transferrin receptor (TfR) and CD98hc, is a transformative approach enabling more effective brain delivery of biotherapeutics for treatment of neurological diseases. TfR-mediated delivery promotes rapid, high brain uptake, while CD98hc-mediated delivery is slower with more prolonged exposure. Here, we engineer a huIgG Fc domain to bind both TfR and CD98hc to create a dual transport vehicle (TV) platform that drives distinct brain delivery properties. Dual TVs achieve significantly higher brain concentrations than TVs targeting either TfR or CD98hc alone. Modulation of TfR and CD98hc affinities shifts dual TV brain exposure kinetics and biodistribution. Stronger TfR affinity drives faster brain uptake and clearance, while stronger CD98hc affinity yields higher, more sustained concentrations, likely due to CD98hc affinity-dependent reduction in TfR-mediated neuronal internalization. This dual targeting strategy leverages the complementary properties of TfR and CD98hc-mediated brain exposure to increase optionality for brain delivery of biotherapeutics.

## Introduction

Delivering biotherapeutics across the blood-brain barrier (BBB) represents a substantial hurdle in successfully treating neurodegenerative diseases such as Alzheimer’s, Parkinson’s, amyotrophic lateral sclerosis, brain enzyme deficiencies, and central nervous system (CNS) tumors and metastases^1–5^, as only a small fraction of systemically administered large molecules typically reach the brain parenchyma^6–8^. Recent studies have suggested the predominant mechanism of brain entry for most traditional protein drugs may be initial transport across blood-CSF barrier sites followed by subsequent transport from CSF into brain parenchyma^9–14^. This transport path results in restricted biodistribution and often insufficient target engagement, severely limiting therapeutic efficacy. Targeting proteins that are highly expressed on brain endothelial cells (BEC) has emerged as a promising strategy for brain delivery of peripherally administered biotherapeutics^3,8,14–27^. The viability of this delivery approach has been recently supported by promising clinical data^28–31^. The most well characterized BBB-targeting proteins include transferrin receptor 1 (TfR)^8,12,14–16,18–20,24,32–40^ and CD98hc^17,23,25,26^. TfR-mediated brain delivery is characterized by high and rapid brain uptake, due to a fast rate of cellular uptake and internalization^41,42^. This also contributes to a relatively fast clearance from brain, likely driven by TfR-mediated internalization into parenchymal cells, including neurons^8,12,35^. Conversely, brain delivery via CD98hc binding exhibits much slower kinetics and is characterized by prolonged retention^17,25^. Experimental evidence suggests that CD98hc-binding molecules predominately localize to the cell surface (including astrocyte processes) and exhibit much slower internalization kinetics compared to TfR^17,25^.

Optimizing large molecule biodistribution for enhanced or novel functionality has been accomplished by targeting additional proteins or epitopes with bispecific antibodies^43,44^. In this study, we investigated if combining TfR and CD98hc binding in a single molecule offers additional benefits for brain uptake, retention, and biodistribution. We utilized the previously reported transport vehicle (TV) platform, which utilizes a modified huIgG Fc domain engineered to bind TfR or CD98hc^8,17^. When combined with knob-into-hole heterodimerization^45^, this format enables monovalent Fc binding to both TfR and CD98hc, while maintaining an antibody-like structure and negating the need for non-native linkers or additional domains. Our findings demonstrate that such a bispecific approach, which we term a dual TV, further increases brain concentrations compared to TVs targeting either TfR or CD98hc alone. By altering the affinity for TfR and CD98hc, we can fine-tune CNS parenchymal cell-type biodistribution as well as modulate brain exposure kinetics toward either higher TfR-mediated uptake or higher CD98hc-mediated retention. This dual TV approach has the potential to enable further optionality for brain delivery of therapeutics by harnessing key properties of both TfR- and CD98hc-mediated brain delivery and CNS biodistribution.

## Results

### Engineering the binding properties of a TfR/CD98hc dual TV

To explore the potential of targeting both TfR and CD98hc for brain delivery, we generated a dual targeting molecule by combining previously characterized TfR-targeting TV35^8^ and CD98hc-binding TV6^17^ families on a human IgG1 (huIgG) antibody Fc region using knob-into-hole technology^45^ (**Figure 1a**). The resulting dual TV consists of an engineered Fc in which one CH3 domain binds human TfR and the other CH3 domain binds human CD98hc. To isolate the effects of the dual TV, we paired the affinity variants TV35.23.4 (K_D_ = ∼620 nM) and TV6.8 (K_D_ = ∼170 nM) with Fabs that lack specific binding to any endogenous target^46^, generating a dual antibody transport vehicle (ATV) that we will refer to as ATV.TfR 620/CD98hc 170. We confirmed that this dual ATV has an affinity for both TfR and CD98hc comparable to the single-binding ATVs, indicating that each binding patch functioned independently without being impacted by the presence of the other (**Table S1**).

**Figure 1:**
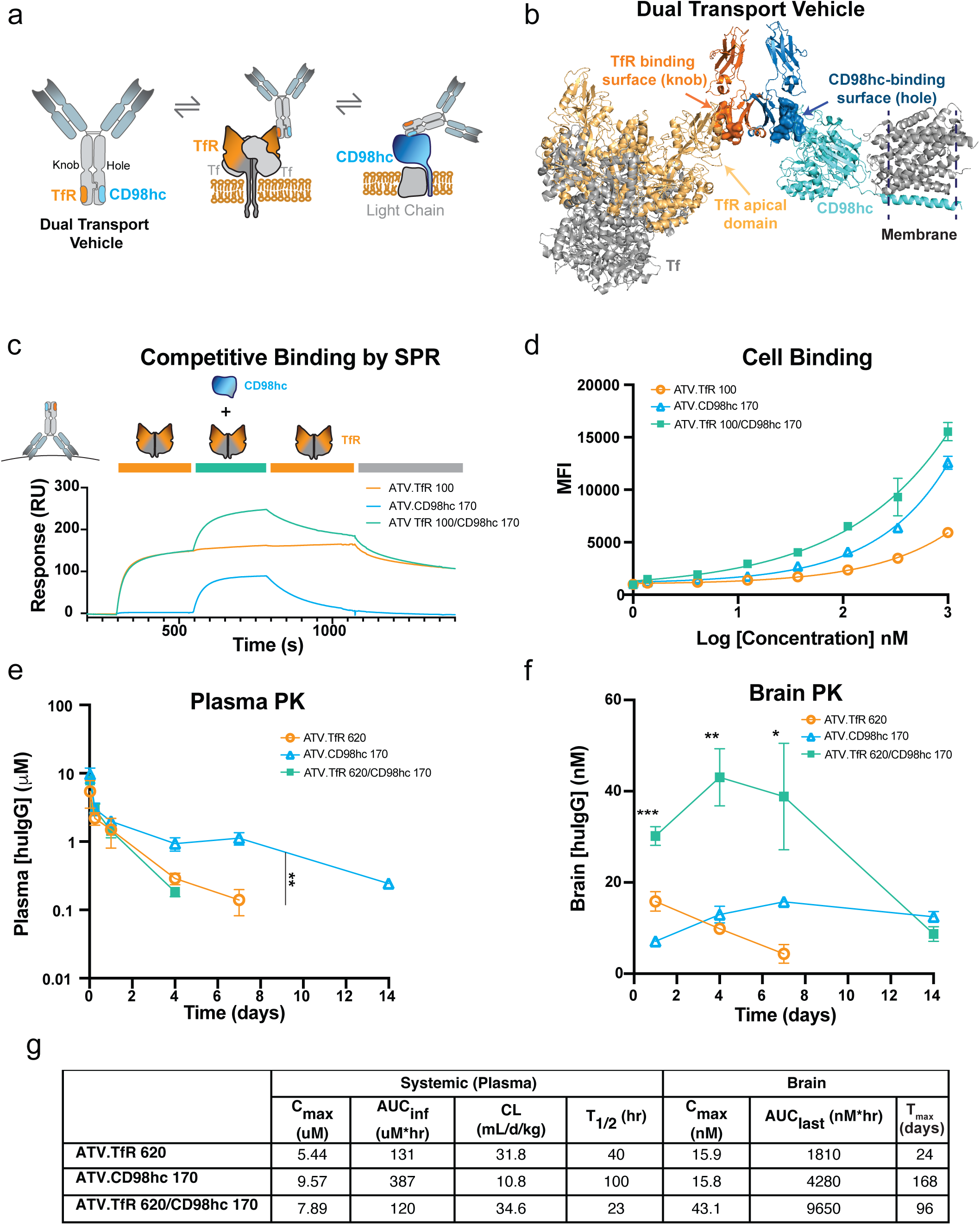
Generation, characterization, and brain exposure of dual TfR and CD98hc binding TVs. **(a)** Cartoon representation of the dual ATV: each Fc strand of huIgG was engineered to bind to TfR or CD98hc (**b**) A model of the TV35:TV6.6 dual TV bound to the full-length TfR^ECD^ plus bound transferrin and CD98hc^ECD^ plus its transmembrane domain binding partner, LAT1, showing a possible orientation of simultaneous binding. **(c)** Binding measured by SPR of 300 nM TfR^ECD^ in the presence or absence of 300 nM CD98hc^ECD^ to immobilized dual ATV.TfR 100/CD98hc 170 (teal), ATV.TfR 100 (orange) or ATV.CD98hc 170 (blue). (**d**) Geometric mean fluorescent intensity (MFI) of ATV.TFR 100, ATV.CD98hc 170, and dual ATV.TfR 100/CD98hc 170 binding on HEK293T cells by flow cytometry. Graph represents mean +/- SD from n= 3 technical replicates **(e-f)** Plasma and brain pharmacokinetics (PK) after a single 25 mg/kg i.v. dose of dual ATV.TfR 620/CD98hc 170 and affinity matched single ATV.TfR and ATV.CD98hc controls. Graphs represent mean ± SD for n=3-4/group per time point (missing values are below LLOQ for the assay). * p < 0.05, **p < 0.01, ***p < 0.001 by mixed effects-model analysis. Exact p values listed in Table S3 (**g**) Non-compartmental analysis of plasma and brain PK.

Molecular modelling using high resolution structures of TV35 bound to a circularly permuted TfR apical domain^8^ and TV6.6 bound to the extracellular domain (ECD) of CD98hc^17^ suggests there is sufficient space for transferrin-bound TfR ECD (TfR^ECD^) and CD98hc ECD (CD98hc^ECD^) to simultaneously bind the dual ATV in solution (**Figure 1b**). To determine if simultaneous binding of TfR and CD98hc by the dual ATV is possible in solution, we measured the affinity for each receptor using surface plasmon resonance (SPR) in the presence of a saturating concentration of the other protein. Under these conditions, binding curves were comparable to those obtained in the absence of the other protein, indicating that neither TfR^ECD^ nor CD98hc^ECD^ competes with the ability of dual ATV to bind the other target and supporting the possibility for simultaneous engagement of both targets in solution (**Figure 1c, S1a-c**). However, based on modeling, simultaneous co-engagement of cell membrane localized TfR and CD98hc could require significant membrane curvature and/or reorientation of the CD98hc^ECD^ or TfR^ECD^ and thus likely restricts the possible geometries required for simultaneous binding. Therefore, to better understand the potential for dual ATVs to simultaneously bind membrane localized TfR and CD98hc, we used flow cytometry to measure binding to HEK293 cells, which express both human CD98hc and TfR. Compared to ATVs that target only TfR or CD98hc alone, dual ATVs exhibited a slight increase in cell surface binding (**Figure 1d**), suggesting that avidity could contribute to the binding property of dual ATV on cells that express both targets. However, the observed increase in cell binding of the dual ATV was relatively small compared to what would be expected from a highly avid interaction, suggesting that simultaneous binding either does not occur frequently on the cell surface, or that TfR and CD98hc are not localized to the same membrane domain to allow for simultaneous dual ATV binding.

### Enhanced brain exposure of dual ATVs

We next sought to characterize the effects of dual TfR and CD98hc binding on plasma and brain exposure. ATV.TfR 620/CD98hc 170 and the corresponding single-binding ATV.TfR 620 and ATV.CD98hc 170 molecules were administered intravenously (i.v.) at 25 mg/kg in TfR^mu/hu^; CD98hc^mu/hu^ double knock-in (DKI) mice. The dual ATV had the highest clearance from plasma (**Figure 1e, g**), likely due to increased target-mediated deposition resulting from binding to both TfR and CD98hc, which are each expressed in the periphery but with different patterns^12,47–49^. Consistent with previous reports^8,17^, ATV.TfR 620 showed rapid brain uptake with a T_max_ at 1 day post dose, followed by a steady clearance; in contrast, ATV.CD98hc 170 exhibited slower uptake with a brain T_max_ at 7 days and maintained exposure out to 14 days post dose (**Figure 1f, g**). Strikingly, the dual ATV exhibited significantly higher overall brain concentrations, with brain C_max_ and AUC more than 2.25 times higher than those achieved with single-binding TfR- or CD98hc-only ATVs (**Figure 1f, g**). Brain T_max_ of the dual ATV occurred at an intermediate time point between the T_max_ observed for ATV.TfR and ATV.CD98hc (7 day)^8,17^, suggesting that dual ATVs exhibit brain exposure kinetics influenced by both TfR and CD98hc binding. Overall, these results demonstrate that dual ATV can significantly increase brain concentrations beyond those achieved with single binding ATVs.

### Modulation of brain and peripheral exposure kinetics with dual ATV

To further explore how modulation of TfR and CD98hc affinities impacts brain delivery, we generated nine unique dual ATVs with a range of TfR and CD98hc affinities (**Table S1**). Plasma (**Figure 2a, c, e**) and brain pharmacokinetics (PK) (**Figure 2b, d, f)** of these dual ATVs were evaluated in the DKI mice, along with representative single-binding controls and a non-binding control huIgG. The plasma PK of dual ATVs with strong binding affinity to TfR (K_D_= 100 nM) were minimally impacted by the presence of CD98hc binding, independent of affinity (**Figure 2a, Table S2**). By contrast, plasma PK of dual ATVs with the weakest TfR affinity (K_D_= 2500 nM) were significantly influenced by varying CD98hc binding affinity (**Figure 2e, Table S2**). This pattern was observed to a lesser extent by molecules with mid-affinity (K_D_= 620 nM) TfR binding (**Figure 2c, Table S2**). These observations suggest that TfR-mediated deposition had a much greater impact on plasma exposure compared to CD98hc binding.

**Figure 2:**
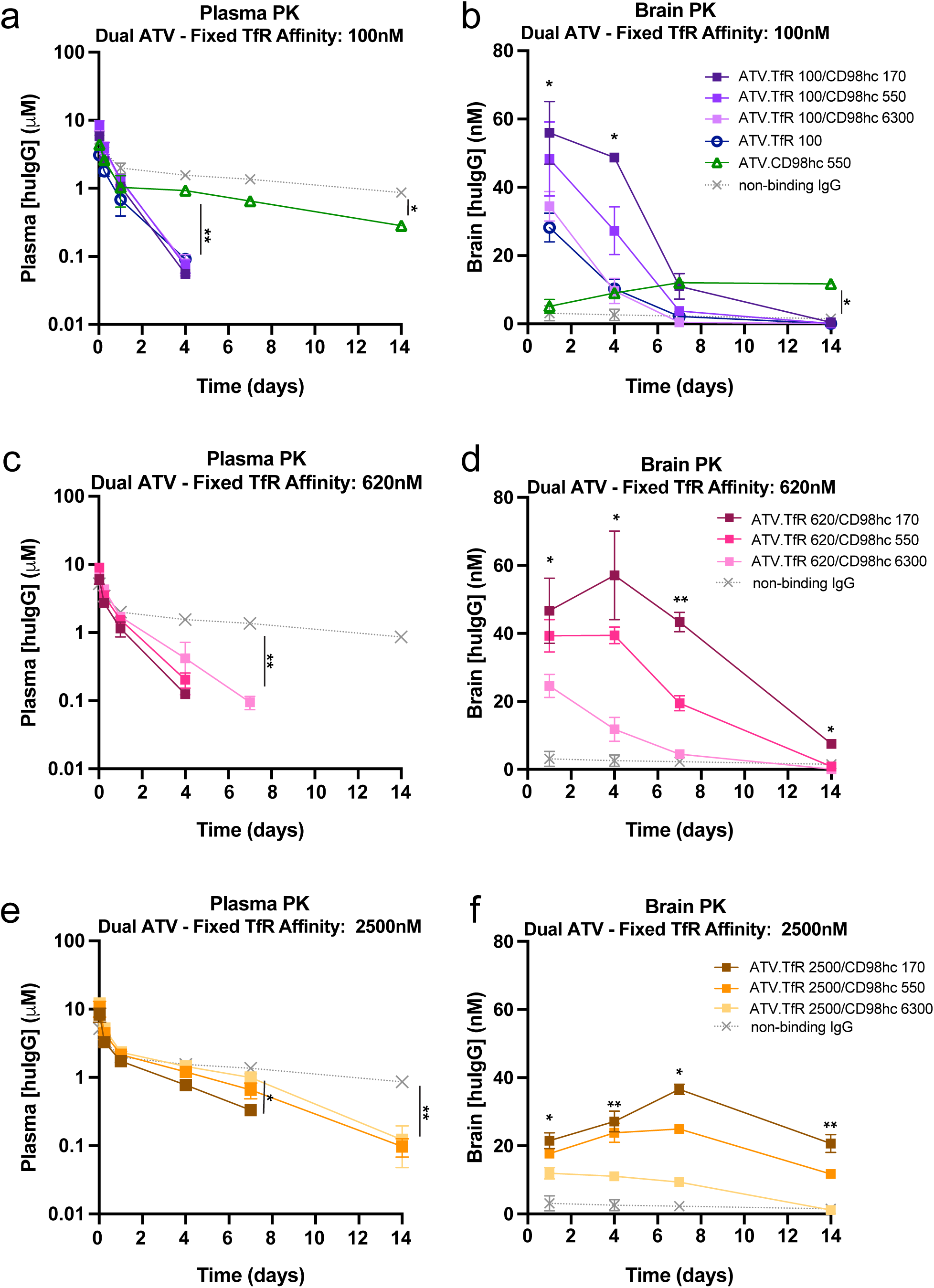
Modulating TfR and CD98hc affinities on dual ATVs enables fine-tuning of brain exposure kinetic profiles. PK analysis of dual ATVs with varying TfR and CD98hc affinities as well as representative TfR and CD98hc single binding ATVs and a non-binding huIgG 25 mg/kg after a single i.v. dose in TfR^mu/hu^/CD98hc^mu/hu^ DKI mice. Plasma and brain huIgG concentrations for dual and single ATVs with TfR affinity of 100 nM (**a-b**), 620nM (**b-c**), and 2500nM (**e-f**), with CD98hc affinities ranging from 170, 550, and 6300 nM (**a-f**). The same non-binding control IgG is shown in each panel for comparison. Graphs represent mean ± SD for n=3-4 mice/group (missing values are below LLOQ for the assay). * p < 0.05, **p < 0.01 by mixed effects-model analysis. Exact p values listed in Table S3.

Dual ATVs with strong TfR binding (K_D_=100nM) resulted in fast brain uptake, with a T_max_ similar to the single binding ATV.TfR at 1 day post dose (**Figure 2b, Table S2**). Brain T_max_ for dual ATVs with weaker TfR affinities was influenced by CD98hc binding and exhibited a right-shifted T_max_ toward 4 and 7 days post-dose (**Figure 2d, f, Table S2**), which was more similar to the single binding ATV.CD98hc (**Figure 2b**). Dual ATVs with stronger CD98hc binding resulted in higher brain concentrations compared to weaker CD98hc variants, along with more prolonged brain exposures (**Figure 2b, d, f, Table S2**). Across all TfR affinities, the brain exposure AUC rank order correlated with CD98hc affinity (**Table S2**). Collectively, these findings indicate that brain exposure magnitude and kinetics can be modulated by adjusting TfR and CD98hc affinities on a dual ATV format.

To investigate how individual TV affinities may influence peripheral organ biodistribution of the dual ATV, we measured huIgG concentrations in bone marrow and kidney (**Figure S2**), which highly express TfR and CD98hc, respectively, and where ATV^TfR^ and ATV^CD98hc^ have been reported to localize^12,47–49^. CD98hc affinity had minimal impact on TfR-mediated bone marrow localization, though strong CD98hc affinity slightly increased exposure **(Figure S2a, c, e**). Consistent with expression and previous biodistribution reports, dual ATVs with strong CD98hc affinity resulted in a higher exposure in the kidney (**Figure S2b, d, f)**. TfR binding at any affinity reduced exposure in the kidney compared to the single-binding CD98hc ATV control, potentially due to shifted biodistribution to bone and/or faster plasma clearance (**Figure S2b, d, f).** These data reveal that bone marrow exposure profiles of dual ATVs are predominantly governed by TfR affinity, while kidney exposure profiles appear to be strongly influenced by CD98hc affinity.

### Modulation of dual ATV cellular internalization and surface retention in vitro

To better understand the cellular mechanisms underlying enhanced dual ATV brain exposure, we assessed whether dual ATVs exhibit both TfR-mediated internalization and CD98hc-mediated surface retention previously described for ATV.TfR and ATV.CD98hc *in vitro*^17^. HEK293T cells were incubated with ATV.TfR, ATV.CD98hc, or dual ATV molecules with varying affinities for 2 hours at 37°C. After incubation, the cells were washed and stained under either permeabilizing or non-permeabilizing conditions to capture total and cell surface only huIgG, respectively (**Figure S3**). Dual ATVs with strong TfR affinity (K_D_= 100 nM) exhibited a similar degree of internalization compared to TfR-only binding ATVs (**Figure S3b**). Compared to TfR-only controls, CD98hc binding increased the cell surface retention of dual ATVs in an affinity-dependent manner **(Figure S3c, e, g**). Notably, dual ATVs with weak TfR affinity (K_D_= 2500 nM) yielded surface retention levels comparable to CD98hc-only binding controls (**Figure S3g**). Together, these results indicate dual ATVs can exhibit both TfR-mediated cellular uptake and CD98hc-mediated cell surface retention and suggest these affinity-dependent trafficking properties may play a key role in the distinct brain exposure kinetics of dual ATVs *in vivo*.

### Brain regional localization of dual ATVs

TfR and CD98hc are expressed by different brain parenchymal cell types, and ATVs targeting each receptor have been reported to have distinct cellular biodistribution patterns^12,17^. To gain insight into the CNS biodistribution of the dual ATV variants, the localization of each was assessed by immunohistochemistry (IHC) at 1 and 7 days post dose, which represent the respective T_max_ of single binding ATV.TfR and ATV.CD98hc. Low magnification images of sagittal sections revealed that the staining intensities of dual and single binding ATVs were overall consistent with huIgG concentration measurements (**Figure 3a, c** compared to **Figure 2b, d, f**). Specifically, dual ATVs with stronger TfR affinity showed brighter staining at 1 day post dose, and dual ATVs with strong CD98hc affinity and weak TfR affinity had brighter staining at 7 days post dose (**Figure 3a, c**).

**Figure 3:**
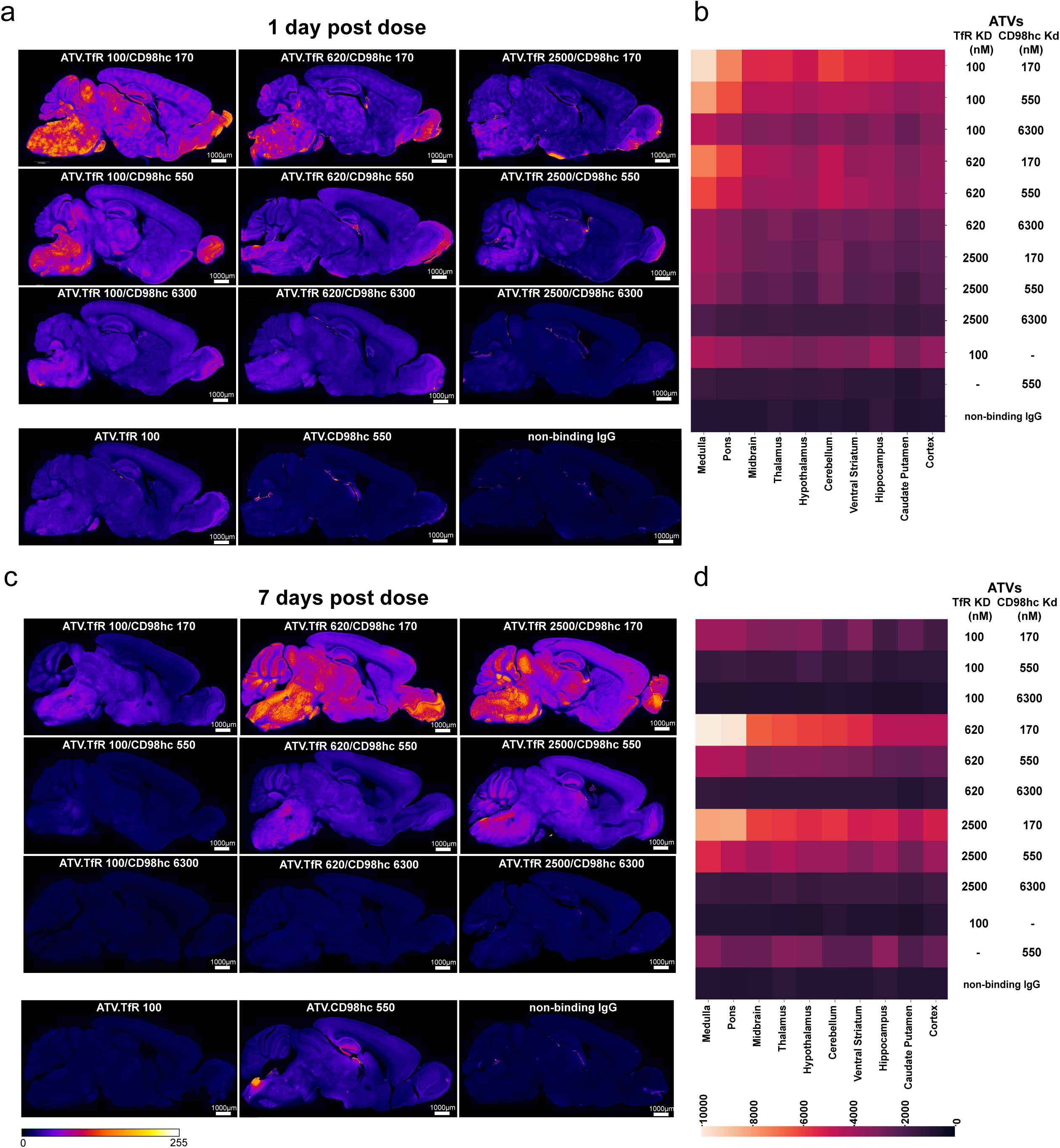
Brain regional localization of dual TV affinity variants. **(a,c)** Immunohistochemical analysis of dual ATV localization in sagittal mouse brain sections as detected with anti-huIgG. Representative images are shown at 1 day post dose (**a**) and 7 days post dose (**c**) following a single 25 mg/kg i.v. administration of dual ATVs. The images illustrate the distribution of dual ATVs with varying TfR and CD98hc affinities across the brain. Single binding ATVs and non-binding huIgG are shown in the bottom panels. Sections were aligned to the Allen brain atlas, ccfv3, sagittal, for regional analysis. (**b, d)** Quantitative heatmaps show brain regional distribution of huIgG at 1 (**b**) and 7 days post-dose (**d**).

Images were registered to the Allen Mouse Brain atlas^50^ to quantify brain regional localization. Consistent with delivery across the BBB, broad distribution of dual ATVs, ATV.TfR, and ATV.CD98hc was observed across multiple brain regions at both time points **(Figure 3b, d)**. Interestingly, staining intensity was proportionally higher for dual ATVs than for ATV.TfR and ATV.CD98hc in brain stem regions (e.g. pons and medulla) at both time points (**Figure 3b-d)**.

In contrast to ATVs and consistent with distinct routes of entry into the brain, it has been previously reported that control IgG is localized almost exclusively to the choroid plexus, brain surfaces leptomeninges, and perivascular compartments^12,14^. Consistent with these previous reports, there was minimal ATV.TfR localization to the leptomeninges and choroid plexus (**Figure S4d, h left panel**), whereas ATV.CD98hc exhibited higher localization to these regions compared to both ATV.TfR and control IgG (**Figures S4d, h; S5d, h**). In the dual ATV format, CD98hc binding increased leptomeningeal and choroid plexus localization in the region of the ambient cistern^10,51,52^ at 1 day post-dose **(Figure S4a-c, S5a-c)**, while mid and strong TfR affinity reduced that localization by 7 days post dose (**Figure S4e-g, S5e-g).** Together, these data suggest that modulating TfR and CD98hc affinities on dual ATVs can shift the CNS biodistribution to be more like ATV.TfR or ATV.CD98hc.

### Decreased neuronal localization of dual ATVs with increased CD98hc affinity

To better understand the cell type-specific biodistribution of dual ATVs, cortical sections were co-stained with aquaporin 4 (AQP4), a marker for astrocyte processes and endfeet^53–55^ shown to colocalize with ATV.CD98hc^17^, and the neuronal marker NeuN, where ATV.TfR has been previously shown to robustly bind and internalize^8,12,25,35^. We additionally used AQP4 staining to aid segmentation of blood vessels, as AQP4 enriched astrocytic end feet prominently surround cerebral blood vessels^5,53–55^. NeuN-positive staining was used to segment neurons, and the remaining tissue area was classified as non-neuronal parenchyma (**Figure 4a-c**, lower right panels).

**Figure 4:**
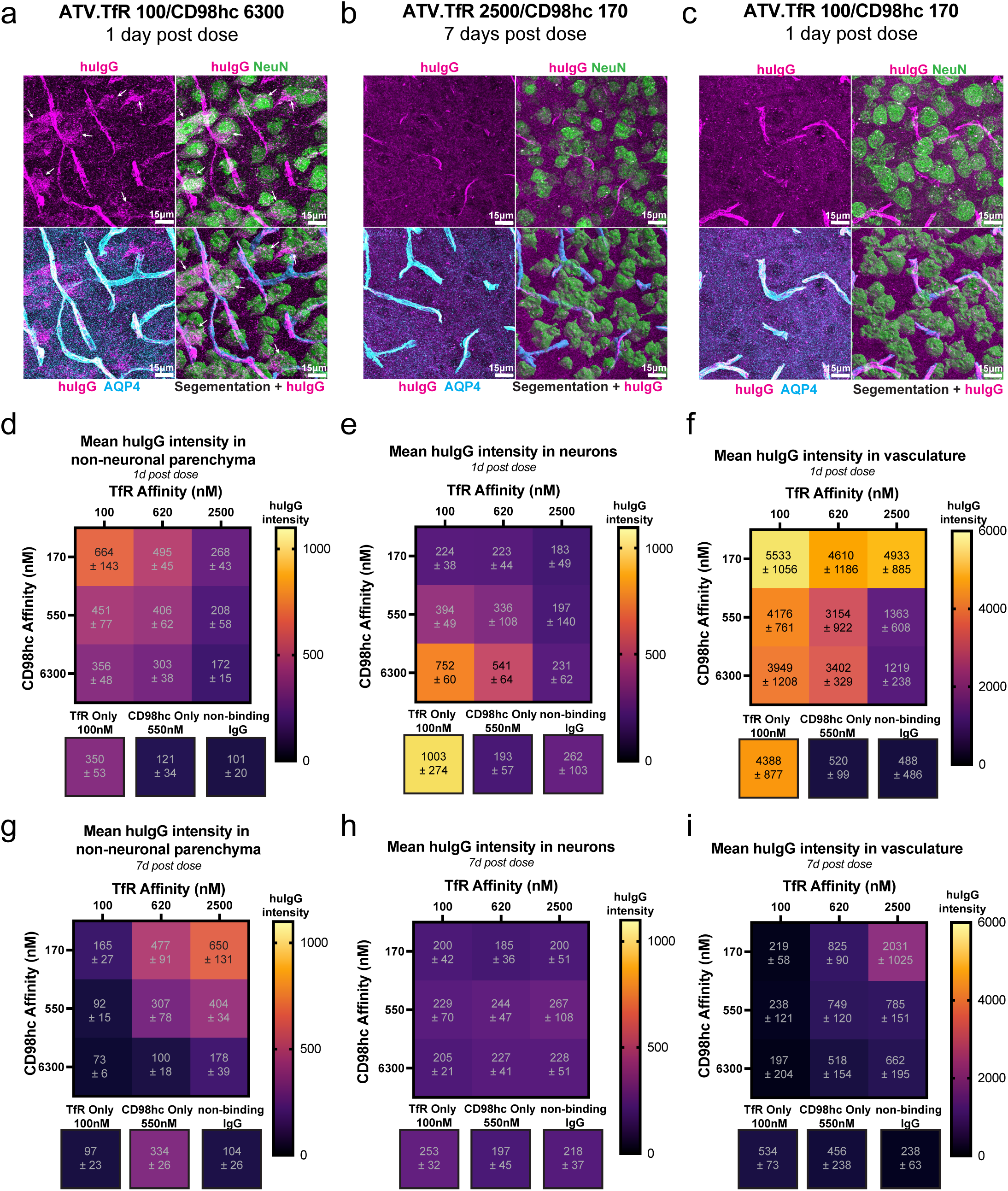
Quantification of brain cell type specific localization of dual ATV affinity variants. **(a-c)** Immunohistochemical analysis showing cell-specific localization of dual ATVs with TfR/CD98hc affinity combinations in mouse brain cortex after a single 25 mg/kg i.v. dose. Representative images show staining for human IgG (huIgG, magenta), NeuN (neuronal marker, cyan) and AQP4 (astrocyte process marker, green). Images were segmented to identify three distinct regions: vasculature (large, bright AQP4 positive structures), neurons (NeuN), and non-neuronal parenchymal (defined as all remaining areas). Representative images are shown for (**a**) ATV.TfR 100/CD98hc 6300 at 1 day post dose, with white arrows pointing at huIgG staining within neurons(**b**) ATV.TfR 2500/CD98hc 170 at 7 days post-dose, and (**c**) ATV.TfR 100/CD98hc 170 at 1 day post dose. **(d-i)** Quantification of huIgG staining intensity in segmented brain regions at 1 (**d-f**) and 7 days (**g-i**) post dose. Heatmaps represent mean intensity levels in non-neuronal parenchyma (**d, g**), neurons (**e, h**), and vasculature (**f, i**) for dual ATV TfR/CD98hc affinity combinations. Values in each square represent mean intensity ± SD. Pearson r = 0.8714, p < 0.001 (see Table S3 for additional details).

HuIgG quantification of the cortex using high-resolution confocal microscopy correlated well with both huIgG intensity in whole sagittal sections by IHC (**Figure S6**, compared to **Figure 3a, c**), as well as bulk brain tissue huIgG concentrations (**Figure 2b, d, f)**. The consistent trends across these orthogonal methods supported the use of cortical IHC images to interrogate the cellular distribution of dual ATVs and gain further insights into the mechanism of dual ATV mediated brain exposure. HuIgG staining intensity in the non-neuronal parenchyma also correlated with total huIgG staining and quantification of huIgG concentrations (**Figure 4d, g**), which is likely due to CD98hc mediated localization to astocytes processes and TfR mediated localization to neuronal processes, which would not be captured by NeuN segmentation which is confined to neuronal cell bodies. Consistent with the reduction in plasma exposure over time, vascular huIgG intensity was highest for all tested variants at 1 day post dose and significantly reduced by day 7 (**Figure 4f, i)**. Consistent with the slower brain uptake kinetics associated with CD98hc binding, some residual vascular huIgG was observed with the dual ATV.TfR 2500/CD98hc 170. Among dual ATV variants, the dual ATV with the strongest TfR binding relative to CD98hc (ATV.TfR 100/CD98hc 6300) had the most prominent neuronal localization at 1 day post dose (**Figure 4a, e**), while the dual ATV with the strongest CD98hc binding relative to TfR (ATV.TfR 2500/CD98hc 170) robustly colocalized with AQP4 within the parenchyma at 7 days post dose (**Figure 4b, g**). Interestingly, at 1 day post dose, CD98hc binding reduced TfR-mediated neuronal localization in a CD98hc affinity-dependent manner; indeed, even weak CD98hc affinity reduced the neuronal localization compared to the single binding ATV.TfR control **(Figure 4e**). Strikingly, the degree of neuronal co-localization for dual ATV.TfR 100/CD98hc 170 was similar to CD98hc-only or non-binding IgG controls at 1 day post dose, suggesting CD98hc binding dramatically reduces neuronal targeting even in the presence of strong TfR affinity (**Figure 4c, e**). The intensity of neuronal huIgG for all tested ATVs was indistinguishable from control IgG at 7 days post dose, suggesting that TfR-mediated neuronal uptake may serve as a clearance pathway for ATVs (**Figure 4h**). Together, these biodistribution data suggest that CD98hc binding reduces TfR-mediated localization and subsequent clearance which prolongs brain exposure observed with dual ATVs.

## Discussion

Here, we demonstrate that engineering a molecule to bind both TfR and CD98hc creates a brain delivery platform with unique brain exposure profiles compared to targeting either receptor alone. The magnitude and kinetics of dual ATV brain exposure are tunable by adjusting the affinities for TfR and CD98hc. Remarkably, a dual ATV with moderate TfR affinity (K_D_= 620 nM) and strong CD98hc affinity (K_D_= 170 nM) achieved more than 2-fold higher brain C_max_ after a single dose compared to single binding ATVs at these affinities (**Figure 1f**). Reducing TfR affinity while maintaining the same CD98hc affinity resulted in sustained brain concentrations above 20 nM for 14 days – an exposure profile not previously achievable with brain delivery platforms that target either TfR or CD98hc alone at this dose level (**Figure 2f)**. In addition to influencing brain exposure kinetics, altering the affinities of TfR and CD98hc also impacted the CNS biodistribution of dual ATVs:CD98hc binding reduced neuronal localization of dual ATVs compared to ATV.TfR, and the magnitude of this effect was dependent on CD98hc affinity (**Figure 4e**).

TfR is known to undergo constitutive ligand-independent recycling from the plasma membrane on the order of minutes^56^, and as a result, molecules that bind TfR enables fast receptor-mediated transport into the brain^8,35–37^. However, the trade-off of this fast cellular internalization is subsequent catabolism, which likely represent a major clearance pathway for TfR-targeting molecules in the brain through TfR-expressing parenchymal cells. On the other hand, CD98hc exhibits much slower internalization kinetics, resulting in both the delayed brain T_max_ of ATV.CD98hc compared to ATV.TfR, as well as longer brain exposure retention previously reported^17^. Thus, dual ATVs takes advantage of both TfR and CD98hc trafficking properties where binding to CD98hc reduces TfR-mediated cellular internalization and enables longer brain retention. This reduction in neuronal localization occurs even with relatively weak CD98hc binding (K_D_= 6300 nM), which may be due to a higher proportion of accessible CD98hc present on the surface of brain parenchymal cells, while a significant fraction of TfR is intracellularly localized^57,58^. Together, these findings support a bispecific approach to create a unique BBB-crossing platform that further enhances overall exposure by reducing brain clearance. A similar effect was recently obtained by combining Fabs to myelin oligodendrocyte glycoprotein (MOG) and TfR to generate a bispecific antibody that resulted in prolonged brain exposure and altered brain biodistribution^59^. Although bispecific molecules are extensively employed in cancer therapeutics to achieve enhanced functionality^43,44,60–63^, these are the first examples of this approach being applied to a brain delivery platform to further optimize performance.

Whether dual ATVs engage both TfR and CD98hc is likely to have a significant impact on their *in vivo* behavior. *In vitro* SPR data and modeling based on crystal structures indicated that dual ATVs can simultaneously bind TfR and CD98hc in solution (**Figure 1b, c, S1**). However, modeling and cell binding data (**Figure 1b, d**) suggested membrane localization of TfR and CD98hc may limit simultaneous binding of dual ATVs on the cell surface. A caveat is this was only investigated in one *in vitro* system, which does not rule out simultaneous binding in other cell types where expression levels and/or localization of each protein may be different, or within intracellular vesicles where membrane curvature may be markedly different than on the cell surface. Therefore, it remains unclear if, and when, simultaneous TfR and CD98hc binding could occur *in vivo*. Given the low probability of a trans-cell interaction, any simultaneous binding *in vivo* would likely be restricted to cells that co-express TfR and CD98hc within the same cellular compartment. Different binding architecture formats with distinct epitopes and geometries may also alter engagement with TfR and CD98hc resulting in potentially different brain exposure profiles.

Tissues and cells known to express both TfR and CD98hc include brain endothelium, kidney tubules epithelial cells, and bone marrow erythroid cells^17,49,64,65^. Consistent with these expression patterns, ATVs targeting TfR or CD98hc have been shown to localize to these tissues^12,17^. For dual ATVs, increasing affinity for both TfR and CD98hc enhanced localization to the brain vasculature (**Figure 4f**), while marginally and variably increasing distribution to the bone marrow (**Figure S2a**) and reducing localization to the kidney (**Figure S2b**). This is likely due to differences in protein densities or subcellular distribution among different tissues and cell types. These results suggest that while it is possible that simultaneous TfR and CD98hc binding occurs in brain endothelial or bone marrow erythroid cells, the contributions of such an effect to biodistribution is minimal. Additionally, if simultaneous binding does occur on the surface of brain endothelial cells *in vivo*, it may not impact brain uptake as much as other factors such as receptor trafficking, internalization kinetics, and expression level. Further research is needed to fully characterize the tissues and cells that co-express these two receptors, including in the brain stem where dual ATVs had a marked increase in localization (**Figure 3b, d**) that may be explained by potentially higher co-expression of TfR and CD98hc compared to other brain regions. Additionally, more work is warranted to explore how dual targeting could affect these co-expressing cells.

Pairing therapeutics with a specific brain delivery platform requires careful consideration of the desired kinetics, distribution for each treatment, and potential combinatorial effects between the therapeutic and brain delivery targets. The high TfR-mediated uptake combined with low neuronal internalization observed with dual ATVs may well benefit certain therapeutic classes, such as certain cell surface receptor antagonists, antibodies targeting extracellular proteins, and/or therapeutic targets requiring high levels of drug exposure for prolonged periods of time. Additional target binding (e.g., Fabs) or therapeutic cargo (e.g., enzymes, proteins, or oligonucleotides) will almost assuredly alter the peripheral and CNS distribution and exposure kinetics of dual TV therapeutics and thus will need to be evaluated on a case-by-case basis. Nonetheless, the dual TV platform represents a unique and significant advance toward additional optionality in brain delivery strategies critical for developing effective treatments for CNS disorders with diverse underlying biology.

## Methods

### Protein design and purification

Recombinant biotinylated CD98hc and TfR were cloned, expressed and purified as previously described for human CD98hc^ECD^ ^17^, human TfR^ECD^, and circularly permuted TfR apical domain^8^. Single target monovalent TV controls for CD98hc^17^ or TfR^8^ were expressed and purified using KiH technology^45^ with the TV patch on the “knob”, effector function attenuating mutations LALAPG and non-binding DNP02 Fabs, as previously^8^. TV affinity variants were engineered from a parent TV designed to bind TfR (TV35.x)^8^ or parent TV to CD98hc (TV6.x)^17^ by combining mutations at positions known to affect target affinity (∼1-4 AA) and selected using yeast surface display^8^ and/or SPR after recombinant expression. Clones were selected that bound TfR (TV35.23.2, TV35.23.4, and TV35.d1.10) or CD98hc (TV6.8, TV6.d6, and TV6.d28. with a range of binding affinities with otherwise similar physical properties. Dual ATVs were constructed by cloning the single CD98hc TV patch onto a “hole” vector backbone and using the existing TfR patch on the “knob” vector. The mammalian expression vectors encoding the Fab light chain, TfR(knob) and CD98hc(hole) heavy chains were combined, expressed and purified from HEK293 cells to generate dual ATVs as with single controls. Dual ATV clones were tested by SPR to have similar affinities as their single target ATVs as referenced in (**Table S1**). Material purity and affinity was confirmed using SEC, LC/MS and SPR.

### Modeling

A model of the TV35:TV6.6 dual TV bound to the full-length TfR^ECD^ and CD98hc^ECD^ was created using a structural alignment of the TVs in the co-complex crystal structures of TV35 with a circularly permuted TfR apical domain^8^ (PDB ID: 6W3H) and TV6.6 with a CD98hc^ECD17^ (PDB ID: 8G0M). The full-length TfR^ECD^ with transferrin was modelled from (PDB ID: 1SUV)^66^, which was structurally aligned to the TfR circularly permuted apical domain. CD98hc^ECD^ with the bound light chain transmembrane receptor, LAT1, was modelling using the cryo EM structure) (PDB ID: 6JMQ)^67^. The models were created in pymol. The approximate location of the membrane bilayer is shown as a dotted line.

### Affinity measurement and simultaneous binding

TV affinities were measured on a Biacore(TM) 8K (Cytiva) by SPR using recombinant circularly permuted TfR apical domain and CD98hc^ECD^, as previously^8,17^. Simultaneous binding measurements by SPR were collected on a Biacore (TM) 8K using a CM5 chip with an anti-human Fab Capture Kit (Cytiva 29234601) to immobilize either single TV controls or dual TV as indicated (TfR K_D_ = 100 nM and CD98hc K_D_ = 170 nM). After immobilization and washing of the various TVs, analyte was flowed using the ABA method with A: either 300 nM TfR^ECD^ or CD98hc^ECD^ and B: 300 nM both TfR^ECD^ and CD98hc^ECD^ then followed by dissociation step with buffer alone as indicated. Flow rates were 30 ul/min.

### Cell binding

As described previously^17^, HEK293 cells (ATCC, CRL-1573) were maintained in DMEM (Gibco™ 11995073) +10%FBS (VWR 89510-188) + 1x Pen/Strep (Gibco 15140122). For flow-based cell staining, cells were detached using cell scraper and resuspended in pre-chilled (4C) FACS buffer containing PBS (pH 7.4), 1% BSA and 1mM EDTA. All the molecules were diluted in PBS alone and added to cells in a 96 well plate. After 1hr incubation, cells were washed once with FACS buffer, followed by 8min incubation with fix buffer containing 4% PFA in PBS. Cells were washed three times and stained for 30min with secondary antibody (1:500, Thermo A-21445) diluted in permeabilization buffer containing 0.3% Triton X in FACS buffer. Cells were washed three times and resuspended in 100ul FACS buffer prior to data collection. BDFACS Canto analyzer was used to collect 10,000 live events and data was analyzed in FlowJo software. Data was collected in triplicates and mean fluorescence intensity of Alexa 647 was reported (**Figure 1d**).

### In vitro cell trafficking

HEK293 cells (ATCC, CRL-1573) were maintained as described above, and plated (30,000 cells/well) on 96 well plates (PhenoPlate, Revvity 6055308). Molecules were diluted in DMEM with supplements mentioned above and applied to cells at 1000 nM for 2h at 37°C. Cells were then washed in HBSS, incubated in DMEM with supplements mentioned above for 0h, 1h, 4h, 20h, or 24h at 37°C, and washed in HBSS before fixation or lysis.

For immunocytochemistry: After the last HBSS wash, cells were fixed in cold 4% PFA and stored at 4oC overnight until staining. Cells were blocked in 5% BSA + 0.03% TritonX100 in PBS, incubated with molecules in blocking solution overnight, washed twice in phosphate-buffered saline (PBS), then incubated with secondary antibody (1:1000, Jackson ImmunoResearch 109545), cell mask (1:10000, Thermo C10046), and DAPI (1:2000, Thermo D1306) in 1% BSA in PBS for 1.5 h.

### Animal Care

All procedures in animals were performed in adherence to ethical regulations and protocols approved by Denali Therapeutic Institutional Animal Care and Use Committee. CD98hc^mu/hu^, TfR^mu/hu^ DKI mice^8,17^ were housed under a 12-hour light/dark cycle and had access to water and standard rodent diet (LabDiet 5LG4, Irradiated) ad libitum.

### Animal Dosing and Blood Collection

Mice were administered test articles via i.v. injection at a dose of 25 mg/kg. In-life blood samples were collected via submental bleeding into EDTA tubes to prevent clotting. Blood samples were centrifuged at 14,000 rpm for 7 minutes to isolate plasma. For terminal collections, animals were deeply anesthetized with an intraperitoneal injection of 2.5% Avertin. Blood was collected via cardiac puncture and processed under the same conditions as the in-life blood samples. Following terminal blood collection, mice were transcardially perfused with cold PBS.

### Tissue Processing

Fresh-frozen brain tissue was homogenized using a Qiagen TissueLyser equipped with a 5 mm steel bead. Homogenization was carried out for 6 minutes at 30 Hz. Tissue was homogenized in lysis buffer containing 1% NP-40 in PBS with protease inhibitors, using a volume 10x the tissue weight.

### Quantification of huIgG concentration by ELISA

As described previously^17^, huIgG concentrations in plasma, brain, bone marrow, and kidney (**Figure 1e, f, Figure 2a-f, and Figure S2**) were quantified using a generic anti-human IgG sandwich-format ELISA. Plates were coated overnight at 4°C with donkey anti-human IgG (Jackson ImmunoResearch, #709-006-098) at 1 μg/mL in a sodium bicarbonate solution (Sigma, #C3041-50CAP) with gentle agitation. After coating, plates were washed three times with wash buffer consisting of PBS + 0.05% Tween 20. Assay standards and plasma samples were diluted in PBS + 0.05% Tween 20 containing 1% bovine serum albumin (BSA). Plasma samples were diluted at 1:2000 and 1:20,000, and brain lysates were diluted 1:2, 1:5, and 1:20. A standard curve was prepared with concentrations ranging from 0.41 to 1,500 ng/mL (0.003 to 10 nM), with the lower limit of quantitation (BLQ) at < 0.3 nM (for brain) and <0.1 nM for plasma. Standards and diluted plasma samples were incubated on the plates for 2 hours at room temperature with agitation. Following incubation, the plates were washed three times with wash buffer. For detection, a goat anti-human IgG antibody (Jackson ImmunoResearch, #109-036-098) was diluted to a final concentration of 0.02 μg/mL in blocking buffer (PBS + 0.05% Tween 20 + 5% BSA). Plates were incubated with the detection antibody for 1 hour at room temperature with gentle agitation. After a final 3 washes with wash buffer, plates were developed using TMB substrate for 5–10 minutes. The reaction was quenched by adding 4N H₂SO₄, and absorbance was read using 450 nm absorbance.

### Quantification of huIgG concentration in brain lysate by MSD

Similar to a protocol described previously^17^, the total test article concentrations in brain lysate samples (**Figure 2b, d, f**) were quantified using a generic anti-human IgG sandwich-format electrochemiluminescence immunoassay (ECLIA) on a Meso Scale Discovery (MSD) platform. Briefly, 1% casein-based PBS blocking buffer (Thermo Scientific, 37528) was added to an MSD GOLD 96-well small-spot streptavidin-coated microtiter plate (Meso Scale Discovery, L45SA) and incubated for approximately 1 hour. Following the plate blocking and wash steps, biotinylated goat anti-human IgG (SouthernBiotech 2049-08) at a working concentration of 0.5 μg/mL was added to coat the assay plate and allowed to incubate for 1-2 hours. Subsequently, test samples were diluted (MRD of 1:100 in 0.5% casein-based PBS assay buffer) and added to the assay plate. Following the 1-2 hour incubation in the capture step, a pre-adsorbed secondary ruthenylated (SULFO-TAG) goat anti-human IgG antibody (Meso Scale Discovery, R32AJ) at a working solution of 0.5 μg/mL was added to the assay plate and incubated for approximately 1 hour. An assay read buffer (1X MSD Read Buffer T, R92TC) was then added to generate the electrochemiluminescence (ECL) assay signal, expressed in ECL units (ECLU). All assay reaction steps were performed at ambient temperature and with shaking on a plate shaker (where appropriate). The brain lysate required an MRD of 50 with a dynamic range of 4.9 – 2500 ng/mL with a 10 standard point curve (serially-diluted at 1:2 plus a blank brain lysate sample. Brain lysate sample concentrations were back-calculated off the assay-specific calibration standard curve, which was fitted with a weighted four-parameter non-linear logistic regression. The sample back-calculated concentrations in ng/mL were subsequently converted to nanomolar (nM) as the final sample results.

### PK calculations

Non-compartmental analysis was estimated using Dotmatics software 5.5 (Boston, Mass) using a non-compartmental approach consistent with the intravenous route of administration. Parameters were estimated using nominal sampling times relative to the start of each administration. Samples that were below the quantitation limit (BQL) were omitted. The model linear up log down was used to calculate exposures. Descriptive statistics (mean and standard deviation) were generated using Dotmatics.

### Mouse brain immunohistochemistry

Similar to a protocol described previously^17^, following transcardial perfusion with PBS, hemi-brains were drop-fixed in 4% paraformaldehyde overnight at 4°C. Sagittal brain sections (40 µm) were prepared using a microtome (MultiBrain® Technology by NeuroScience Associates). Brain sections were blocked in a solution containing 5% BSA and 0.3% Triton X-100 for 1–3 hours at room temperature. After blocking, sections were incubated overnight at 4°C with the following primary antibodies, diluted in 5% BSA + 0.3% Triton X-100 (dilution buffer): anti-AQP4 (Millipore, AB2218, 1:500), anti-NeuN (Millipore, MAB377, 1:500), and anti-human IgG-647 (Jackson ImmunoResearch, 709-606-149, 1:500). Following primary antibody incubation, sections were washed three times with PBS and then incubated overnight at 4°C with the following secondary antibodies in dilution buffer: donkey anti-rabbit Alexa Fluor 488 (Invitrogen, A21206, 1:500) and goat anti-mouse IgG1 Alexa Fluor-568 (Invitrogen, A-21124, 1:500). Sections were then washed three times with PBS and mounted in ProLong Glass antifade mounting media (ThermoFisher, P36984).

### Tissue Section Imaging and Analysis

Tissues were imaged with a Zeiss AxioScan Z1 with a 20x air objective (0.8 NA). Tissue sections were aligned to the Allen Brain Atlas using ITK-elastix v0.18^68^ by progressively registering the Nissl-stained atlas image to each section with a 3-stage pipeline of a rigid transform, followed by an affine transform, and then a B-spline transform. The default elastix parameters were used for the rigid and affine stages. The B-spline transform was modified to use a final grid spacing of 400 μm, with a 4-stage fitting pyramid of 16x, 8x, 4x, and then 4x down sampling. Brain regions from the atlas were merged as described in^12^ and mean intensity of huIgG was quantified for each region. The affine aligned sections were warped to the Allen Mouse Brain common coordinate framework (CCFv3)^50^ and anatomically equivalent regions containing the hippocampus and the cerebellum were extracted from each image and co-registered as a virtual stack in Fiji with the ‘Register Virtual Stack’ Plugin using the Rigid feature extraction and registration models^69^.

### Confocal Imaging and Analysis

For each tissue section, a small region of the cortex directly superior to the hippocampus was imaged using a Leica SP8 scanning confocal with a 25x water objective (0.95 NA) followed by Lightning super resolution post processing with default settings for the “global” algorithm. Each image was then down sampled 4x in x and y to reduce processing time, and then the background for each channel was removed by smoothing the image with a gaussian filter with a standard deviation of 14 μm and then subtracting the smoothed image from the original image using the functions in the ndimage module in scipy v1.14^70^. Neuronal cell bodies and vasculature were segmented by thresholding the NeuN and AQPN4 channels respectively, followed by size filtering to remove segments smaller than 250 μm^3^, then mean huIgG intensity was quantified in the neuron, vessel, and non-neuronal parenchyma compartments. Images and segmentations were visualized using napari v0.5.1^71^.

## Contributions

**Conceptualization**: Robert C. Wells, Darren Chan, Gerald Maxwell Cherf, Mark S. Dennis, Robert G. Thorne, Mihalis S. Kariolis, Y. Joy Yu Zuchero, Kylie S. Chew

**Investigation:** Robert C. Wells, Padma Akkapeddi, Roni Chau, Johann Chow, Ellen Chien, Allisa Clemens, Michelle E. Pizzo, Ahlam N. Qerqez, Tiffany Tran, Isabel Becerra, Jason C. Dugas, Joseph Duque, Timothy K. Earr, David Huynh, Do Jin Kim, Amy Wing-Sze Leung, Eric Liang, Hoang N. Nguyen, Yaneth Robles-Colmenares, Holly Kane, Elysia Roche, Patricia Sacayon, Kaitlin Xa, Kylie S. Chew

**Methodology:** Robert C. Wells, Padma Akkapeddi, David Joy, Darren Chan, Arash Moshkforoush, Gerald Maxwell Cherf, Michelle E. Pizzo, Kylie S. Chew

**Formal analysis**: Robert C. Wells, Padma Akkapeddi, Darren Chan, David Joy, Johann Chow, Jason C. Dugas, Kylie S. Chew

**Supervision:** Robert C. Wells, Jason C. Dugas, Joseph Duque, David Huynh, Hilda Solanoy, Mark S. Dennis, Joseph W. Lewcock, Ryan J. Watts, Mihalis S. Kariolis, Robert G. Thorne, Christopher M. Koth, Meredith E. K. Calvert, Y. Joy Yu Zuchero, Kylie S. Chew

**Visualization:** Robert C. Wells, David Joy, Meredith E. K. Calvert, Kylie S. Chew,

**Writing – original draft:** Robert C. Wells, Padma Akkapeddi, David Joy, Johann Chow, Christopher M. Koth, Meredith E. K. Calvert, Y. Joy Yu Zuchero, Kylie S. Chew

**Writing – review & editing:** Robert C. Wells, Padma Akkapeddi, Darren Chan, David Joy, Christopher M. Koth, Robert G. Thorne, Meredith E. K. Calvert, Y. Joy Yu Zuchero, Kylie S. Chew

## Competing interests

R.C.W., P.A., D.C., D.J., R.C., J.C., A.M., G.M.C., E.C., A.C., M.E.P., A.N.Q., T.T., I.B., J.C.D., J.D., T.K.E., D.H., D.J.K., A.W.S.L, E.L., H.N.N., Y.R.C., H.K., E.R., P.S., H.L.T., K.X., H.S., M.S.D., J.W.L., R.J.W., M.S.K., R.G.T., C.M.K., M.E.K.C., Y.J.Y.Z, and K.S.C. are or were paid employees of Denali Therapeutics, Inc. Denali has filed patent application no. PCT/US2024/034457 related to the subject matter of this paper, which includes the discovery and application of dual TVs. R.C.W., P.A., D.C., A.M., G.M.C., M.S.D., M.S.K., R.G.T., Y.J.Y.Z, and K.S.C. are inventors of this patent application. There are no other competing interests.

## Acknowledgements

We would like to acknowledge Butch Benitez, Kevin Rebadulla, and Dominic Sobrepena for their dedicated care of the animals at Denali Therapeutics. The trademarks ATV, TV, and TransportVehicle, whether registered or unregistered, are all trademarks of Denali Therapeutics Inc.

## Supplemental Data and Legends

**Figure S1:**
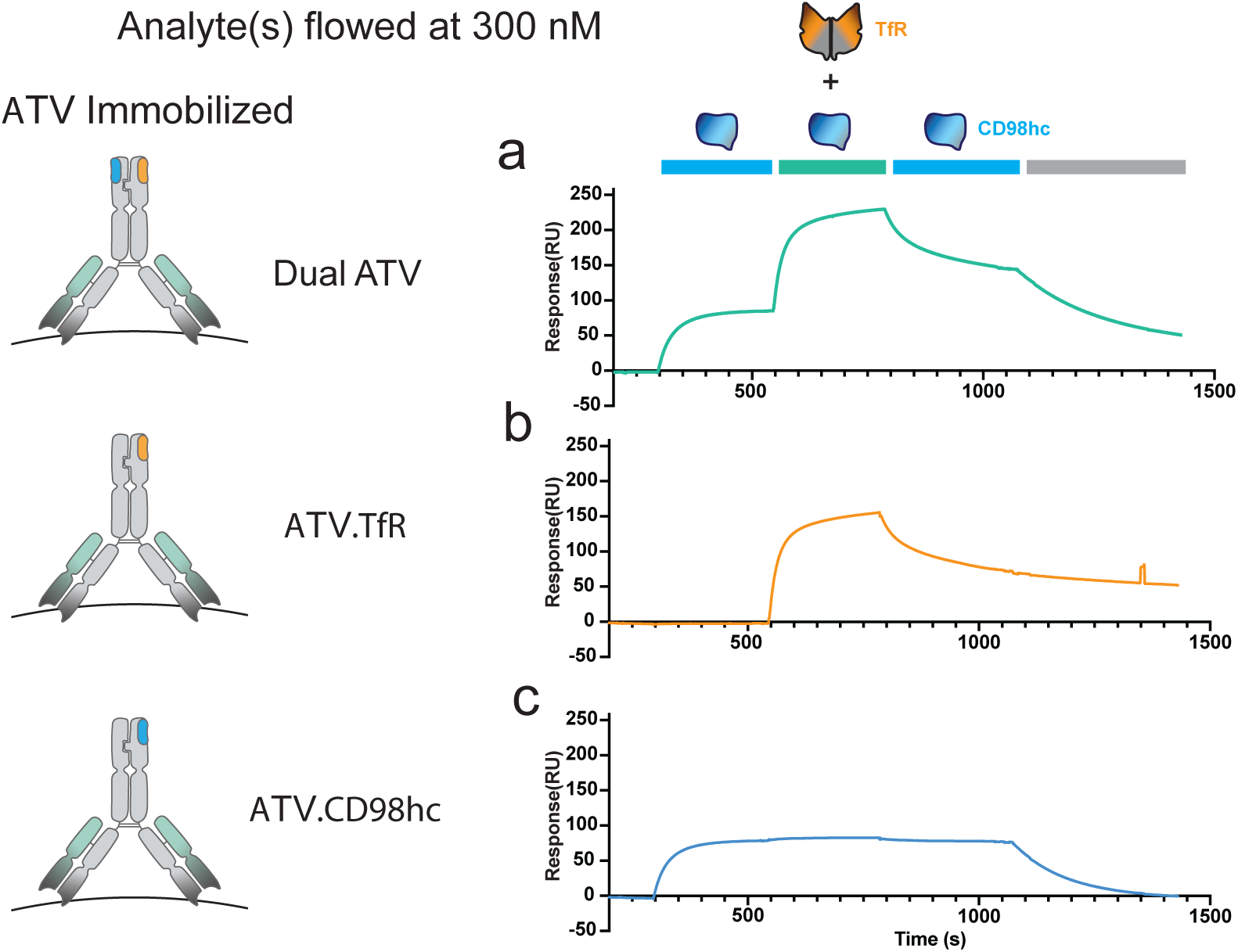
Kinetics of simultaneous binding of dual ATVs and single ATV controls to TfR in the presence of CD98hc by SPR. Dual ATV **(a)**, or the single ATV controls ATV.TfR **(b)** or ATV.CD98hc **(c)** were immobilized on an SPR sensor chip using anti-kappa antibody to ∼150 RU, then sequentially flowed over CD98hc alone, both TfR and CD98hc, CD98hc alone, then buffer alone. The affinity to ATV.TfR (K_D_ = 100 nM) and ATV.CD98hc (K_D_ = 170 nM) were matched to the dual ATV. The ligands were flowed over as indicated above by bars and cartoons at a concentration of 300 nM.

**Figure S2:**
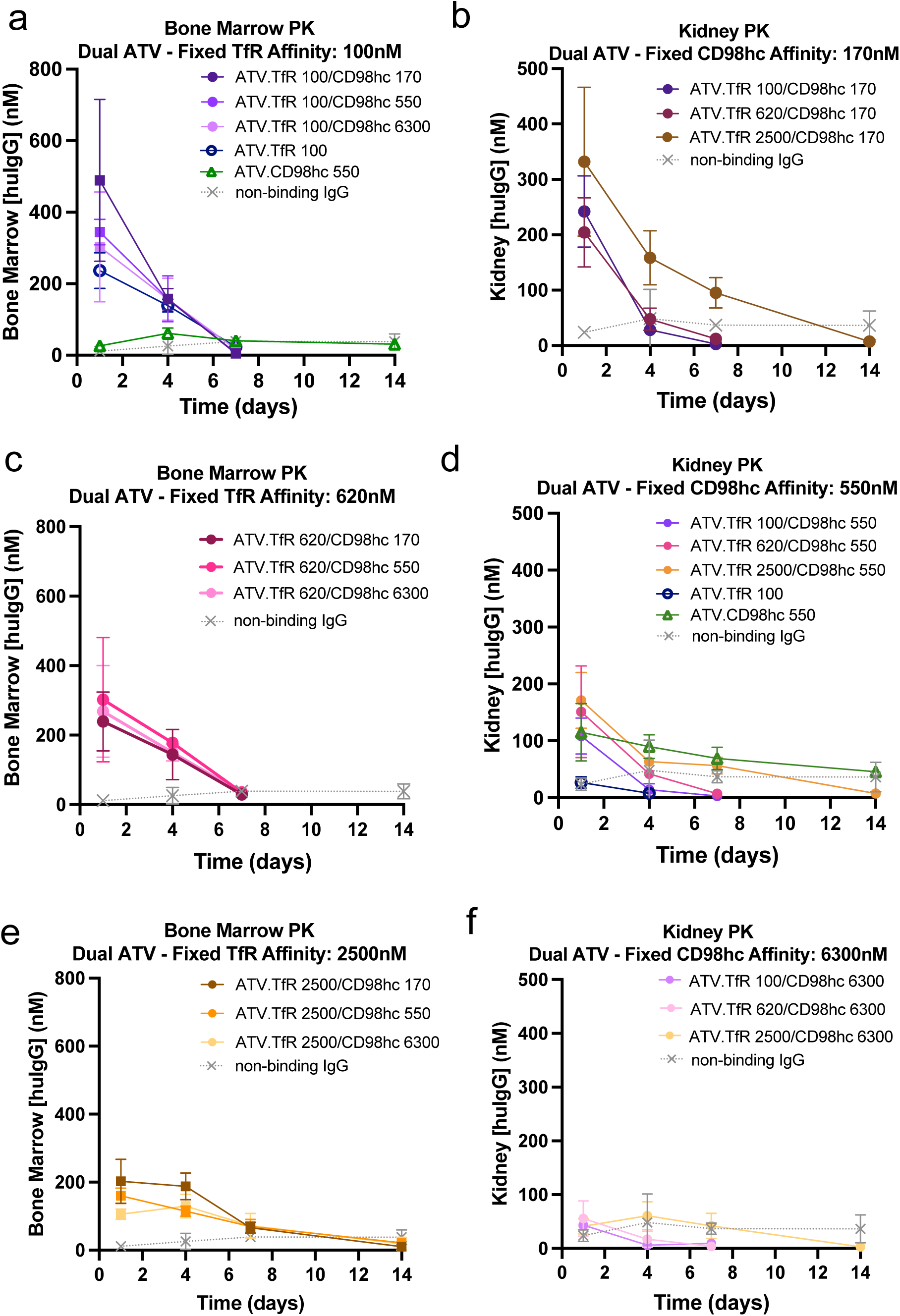
Pharmacokinetics of dual ATVs in bone marrow and kidney. (**a-f**) PK analysis of dual ATVs in bone marrow and kidney following a single 25 mg/kg i.v. dose in TfR^mu/hu^/CD98hc^mu/hu^ DKI mice. (**a, c, e**) Bone marrow PK for dual ATVs with fixed TfR affinities and varying CD98hc affinities (K_D_= 170, 550, and 6300 nM). Kidney PK for dual ATVs with fixed CD98hc affinities and varying TfR affinities (K_D_= 100, 620, and 2500 nM). The non-binding control IgG is shown in each panel for comparison. Graphs represent mean ± SD for n=3-4/group (missing values are below LLOQ for the assay).

**Figure S3:**
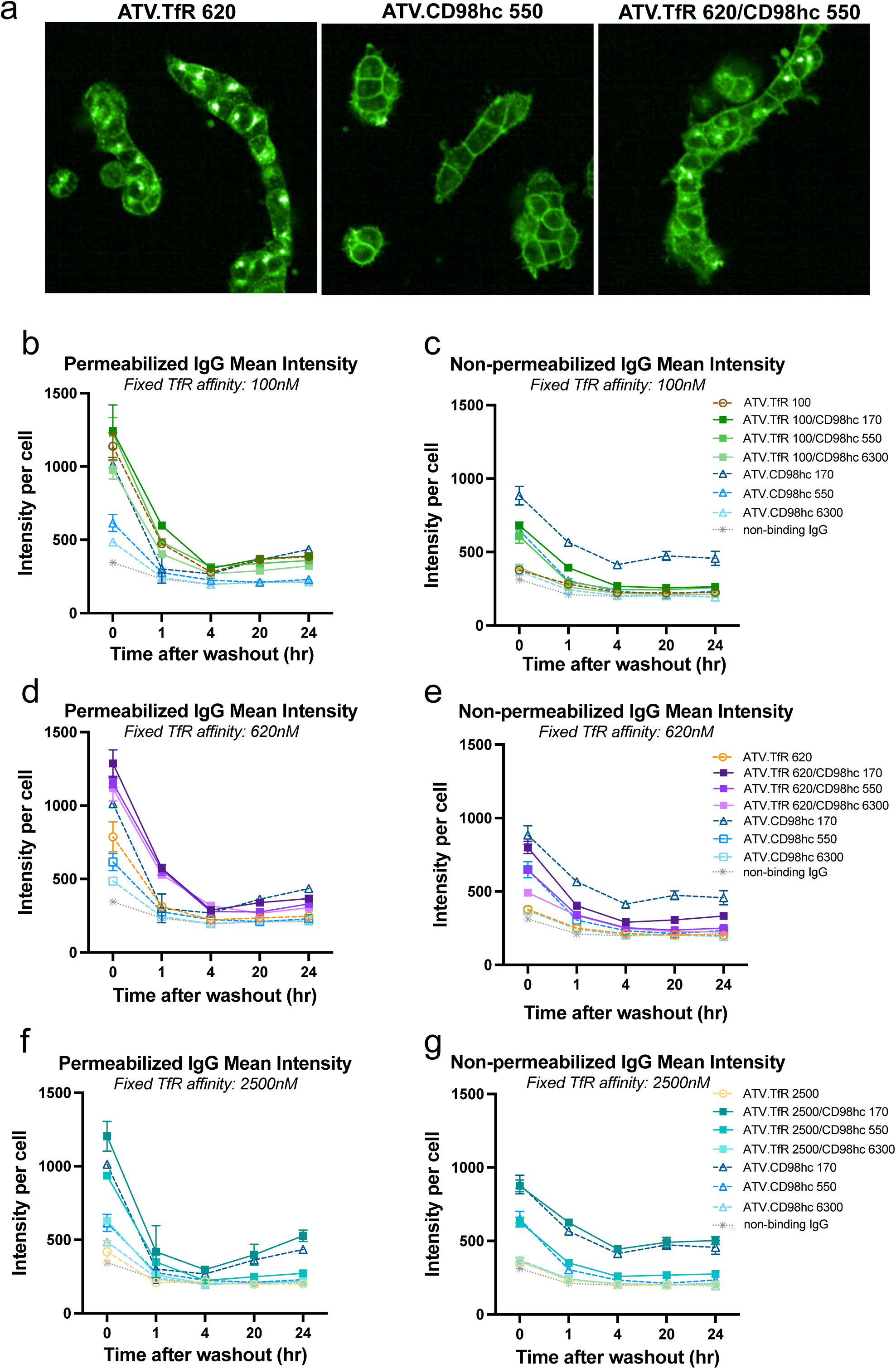
Immunocytochemical analysis of dual ATV cell binding in vitro. **(a)** Representative images of immunocytochemistry showing the binding patterns of a representative dual ATV (right) and its affinity matched TfR-only ATV (K_D_= 620 nM, left) and CD98hc-only ATV (K_D_= 550 nM, middle) under permeabilizing conditions. **(b-g)** Dual ATVs with varying TfR and CD98hc affinities were incubated on HEK293T cells for 2 hours prior to wash out, and immunofluorescence was measured at the indicated time points after incubation at 37°C. Graphs represent mean huIgG intensity per cell +/- SEM, n=2 wells per condition.

**Figure S4:**
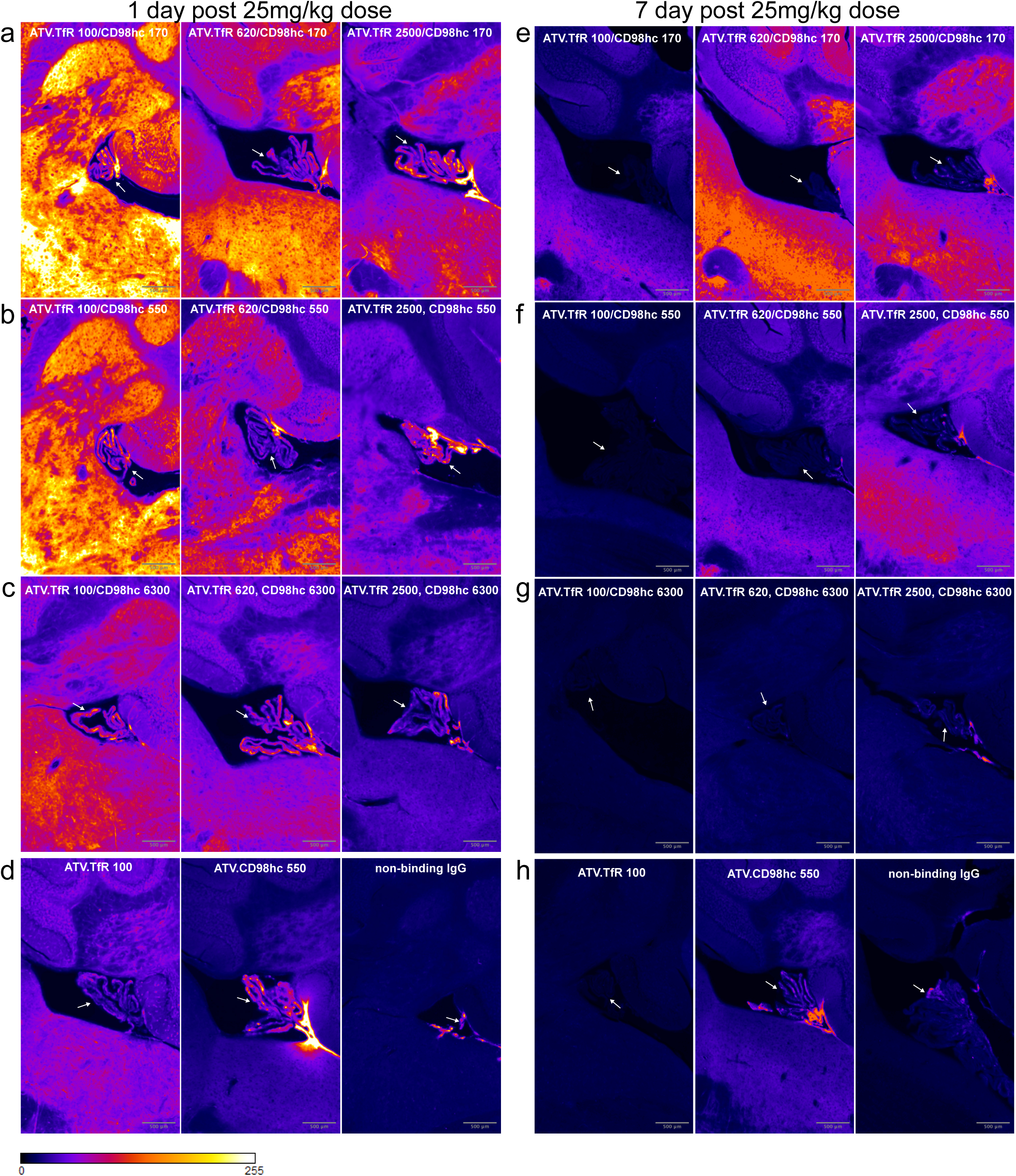
Representative images of dual ATV localization in mouse cerebellum. Representative anatomically equivalent images of dual ATV localization in mouse cerebellum. Hippocampal regions were selected from images of whole sagittal sections (Figure 3), registered to the Allen brain atlas, ccfv3, sagittal, and co-registered as a virtual stack. Arrows indicate the choroid plexus in the 4^th^ ventricle.

**Figure S5:**
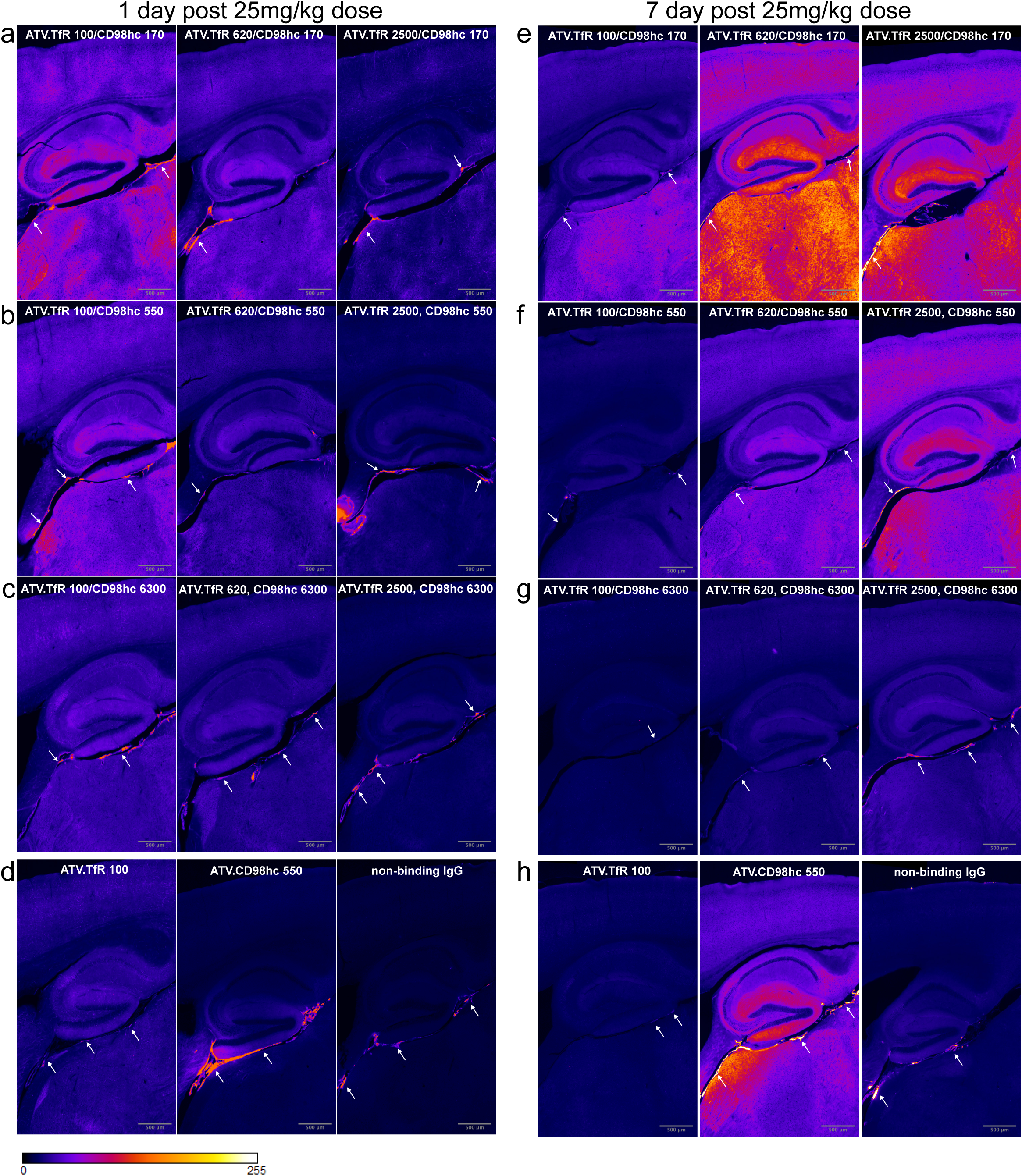
Representative images of dual ATV localization in mouse hippocampus. Representative anatomically equivalent images of dual ATV localization in mouse hippocampus. Hippocampal regions were selected from images of whole sagittal sections (Figure 3), registered to the Allen brain atlas, ccfv3, sagittal, and co-registered as a virtual stack. Arrows indicate leptomeninges in the region of the ambient cistern.

**Figure S6:**
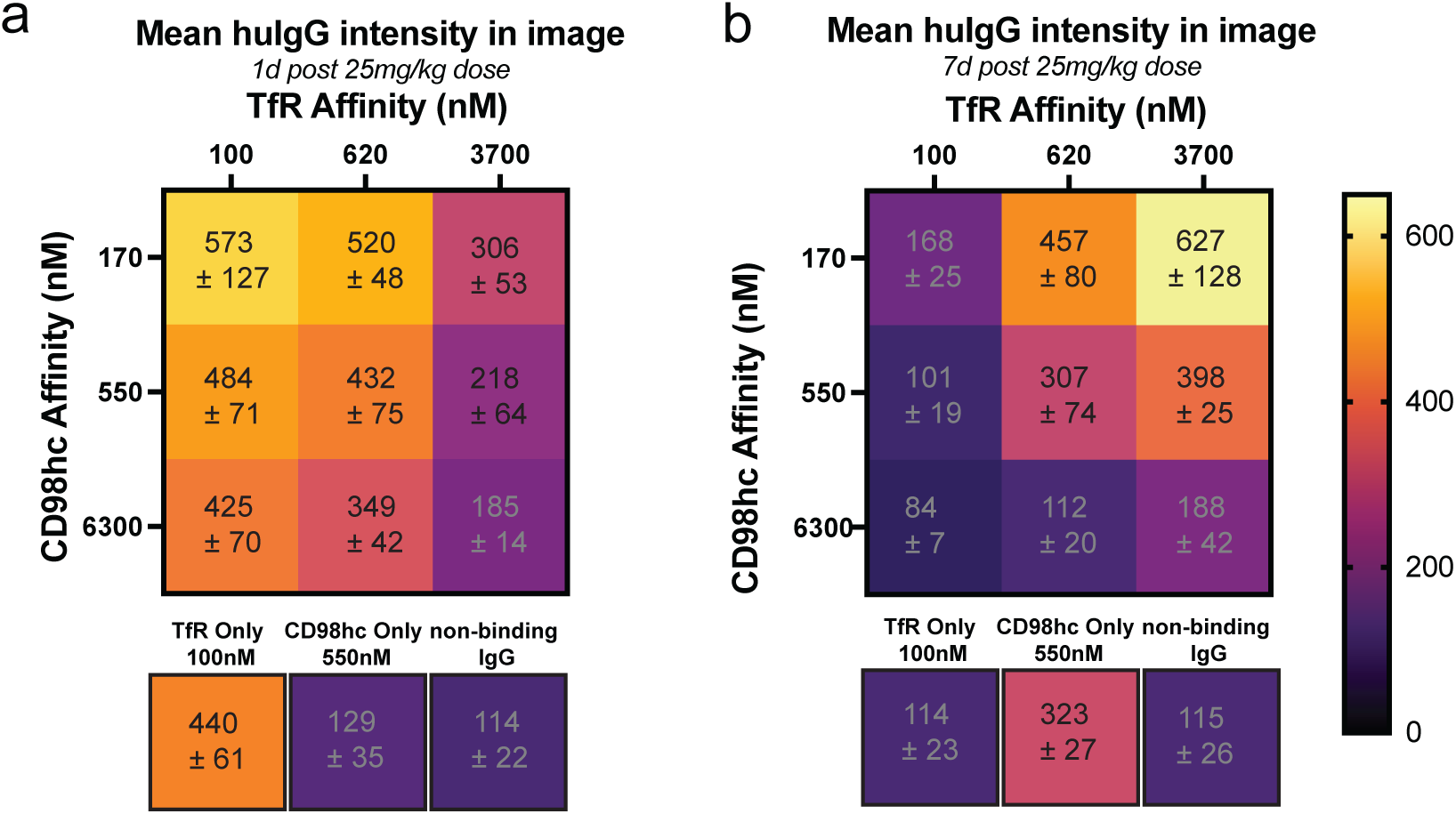
Total huIgG staining intensity for duals ATV in mouse brain cortex. (**a-b**) Quantification of huIgG intensity by immunohistochemistry of dual ATVs with various TfR/CD98hc affinity combinations in mouse brain cortex after a single 25 mg/kg i.v. injection at 1 (**a**) and 7 days post dose (**b**).

---

**Table S1:** Clone names and affinity of single and dual ATVs. Dual ATV names used in the text and figures are matched with the identity of the clone names for TfR and CD98hc TVs. Dual ATVs or single binding control binding affinities to a circularly permuted TfR apical domain and CD98hc^ECD^ (actual) The expected affinities were measured as the average affinity in a single monovalent TV format. Affinities were measured for single and dual ATVs by SPR as described in the methods.

| Short Name | TfR TV clone | CD98hc TV clone | Fab target | Other Fc mutations | Human TfR (apical domain) K<sub>D</sub> (nM) |  | Human CD98hc ECD K<sub>D</sub> (nM) |  |
| --- | --- | --- | --- | --- | --- | --- | --- | --- |
|  |  |  |  |  | Expected | Actual | Expected | Actual |
| ATV.TfR 100/CD98hc 170 | TV35.23.2 | TV6.8 | DNP | LALAPG | 100 | 200 | 170 | 150 |
| ATV.TfR 100/CD98hc 550 | TV35.23.2 | TV6.d6 | DNP | LALAPG | 100 | 200 | 550 | 820 |
| ATV.TfR 100/CD98hc 6300 | TV35.23.2 | TV6.d28 | DNP | LALAPG | 100 | 200 | 6300 | 7400 |
| ATV.TfR 620/CD98hc 170 | TV35.23.4 | TV6.8 | DNP | LALAPG | 620 | 830 | 170 | 150 |
| ATV.TfR 620/CD98hc 550 | TV35.23.4 | TV6.d6 | DNP | LALAPG | 620 | 750 | 550 | 680 |
| ATV.TfR 620/CD98hc 6300 | TV35.23.4 | TV6.d28 | DNP | LALAPG | 620 | 810 | 6300 | 6600 |
| ATV.TfR 2500/CD98hc 170 | TV35.d1.10 | TV6.8 | DNP | LALAPG | 2500 | 5100 | 170 | 130 |
| ATV.TfR 2500/CD98hc 550 | TV35.d1.10 | TV6.d6 | DNP | LALAPG | 2500 | 4900 | 550 | 780 |
| ATV.TfR 2500/CD98hc 6300 | TV35.d1.10 | TV6.d28 | DNP | LALAPG | 2500 | 6000 | 6300 | 5700 |
| ATV.TfR 100 | TV35.23.2 | - | DNP | LALAPG | 100 | 180 | NB | NB |
| ATV.CD98hc 550 | - | TV6.d6 | DNP | LALAPG | NB | NB | 550 | 940 |

NB= no binding

**Table S2:** Non-compartmental analysis of ATV plasma and brain PK.

| ATV variant name | Plasma |  |  |  | Brain |  |  |
| --- | --- | --- | --- | --- | --- | --- | --- |
|  | C<sub>max</sub><br>(uM) | AUC<sub>inf</sub><br>(uM*hr) | CL<br>(mL/d/kg) | T<sub>1/2</sub> (hr) | C<sub>max</sub> (nM) | AUC<sub>last</sub> (nM*hr) | T<sub>max</sub><br>(hr) |
| ATV.TfR 100/CD98hc 170 | 5.83 | 81.0 | 51.4 | 16.5 | 56.0 | 7510 | 24 |
| ATV.TfR 100/CD98hc 550 | 8.48 | 113 | 36.7 | 16.0 | 48.3 | 4780 | 24 |
| ATV.TfR 100/CD98hc 6300 | 8.39 | 110 | 37.9 | 16.6 | 34.4 | 2640 | 24 |
| ATV.TfR 620/CD98hc 170 | 6.10 | 98.1 | 42.5 | 20.7 | 57.1 | 11900 | 96 |
| ATV.TfR 620/CD98hc 550 | 8.98 | 131 | 31.8 | 22.9 | 39.4 | 6810 | 96 |
| ATV.TfR 620/CD98hc 6300 | 8.45 | 175 | 23.8 | 34.5 | 24.6 | 2470 | 24 |
| ATV.TfR 2500/CD98hc 170 | 8.40 | 230 | 18.1 | 60.4 | 36.6 | 9230 | 168 |
| ATV.TfR 2500/CD98hc 550 | 10.4 | 342 | 12.2 | 65.1 | 25.0 | 6600 | 168 |
| ATV.TfR 2500/CD98hc 6300 | 12.1 | 418 | 9.96 | 64.7 | 12.0 | 2420 | 24 |
| ATV.TfR 100 | 3.08 | 59.5 | 70.1 | 21.9 | 28.2 | 2460 | 24 |
| ATV.CD98hc 550 | 4.39 | 310 | 13.4 | 140 | 12.1 | 3380 | 168 |
| Non-binding IgG | 5.20 | 821 | 5.08 | 266 | 2.30 | 652 | 168 |

